# Induction of expansion of ex vivo NK cells using a new feeder cell built in Brazil. An extended flow cytometry evaluation of K562.mbIL21.4BBL

**DOI:** 10.1101/2024.02.23.581719

**Authors:** Caroline Mitiká Watanabe, Caroline Ishihama Suzuki, Alessandro Marins dos Santos, Thiago Pinheiro Arrais Aloia, Grace Lee, David Wald, Oswaldo Keith Okamoto, Julia T. Cottas de Azevedo, Juliana Aparecida Preto de Godoy, Fabio P. S. Santos, Ricardo Weinlich, Lucila N. Kerbauy, Jose Mauro Kutner, Raquel de Melo Alves Paiva, Nelson Hamerschlak

**Affiliations:** Experimental Research Laboratory, Hospital Israelita Albert Einstein, São Paulo, Brazil; Department of Hemotherapy and Cellular Therapy, Hospital Israelita Albert Einstein, São Paulo, Brazil; Human Genome and Stem Cell Research Center, Department of Genetics and Evolutionary Biology, Biosciences Institute, University of São Paulo (USP), São Paulo, Brazil; Oncology and Hematology Center “Familia Dayan-Daycoval”, Hospital Israelita Albert Einstein, São Paulo, Brazil; Department of Pathology, Case Western Reserve University, Cleveland, Ohio USA; Department of Pathology, University Hospitals Cleveland Medical Center, Cleveland, Ohio USA

**Keywords:** NK cells, feeder cells, cytotoxicity, immunophenotyping, immunotherapy

## Abstract

**Background:** Natural killer (NK) cells are lymphocytes from the innate immune system capable of promoting an antitumor response through activation and inhibition receptors. NK cells can better mediate cell engraftment after hematopoietic stem cell transplantation (HSCT), decreasing relapse rates and preventing viral diseases after HSCT. These characteristics make NK cells eligible for application in cell therapy by increasing the frequency of NK cells in peripheral blood. Manufacturing and using feeder cells is necessary to expand NK cells while keeping their cytotoxic characteristics.

**Aims:** We aimed to develop feeder cells from a K562 leukemic cell line capable of promoting clonal expansion of NK cells ex vivo while preserving its antitumor potential.

**Methods and results:** Feeder cells, named K562.clone1, were produced by transduction of mbIL-21 and 4-1BBL proteins. Next, peripheral blood NK cells were co-cultivated with K562.clone1 and expanded more than 100 folds compared to NK cells co-cultivated with K562-WT (less than 10 folds). On the first day of cultivation, the average frequency of NK cells (CD3-CD56+CD16+/−) was 5.95% ± 3.92%, increasing to 83.97% ± 8.19% on the fourteenth day after co-cultivation with K562.Clone1, while the percentage of NK cells raised to 75.15% ± 7.57% when co-cultivated with control. In addition, NK cells expanded with the feeder K562.Clone1 were potentially cytotoxic against acute myeloid leukemia (AML) blast, tumor cell lines of leukemia and glial origin.

**Conclusion:** We successfully built a national feeder cell, named K562.Clone1. The co-culture with K562.Clone1 feeder cell, preserving their primordial functions such as missing-self, important to distinguish health and defective cell, and natural cytotoxicity against tumoral cells, and increasing the natural cytotoxicity.

## INTRODUCTION

Advanced therapies are continually being developed for the treatment of cancer, infectious and genetic diseases since the beginning of cell therapy in 1957, with the first allogeneic transplant performed by E. Donnall Thomas [1]. Cell therapy has evolved and expanded worldwide, becoming the standard procedure for several types of leukemia. Bone marrow transplantation (BMT) is regularly performed in hundreds of centers spread around the world with excellence, thanks to the popularization of protocols, supplies, knowledge, and connectivity. Likewise, cell therapy has broken paradigms, offering a therapeutic option to compassed patients. Moreover, immune cells are being used together in BMT protocols to improve the graft versus host response, favoring a long-term response against leukemia[2, 3].

However, the cost of immunotherapy is a challenge for patients. T cell therapy, approved by the Food and Drug Administration (FDA) in 2017, is one example. The costs of the CAR-T product from Kymriah was initially priced at $475,000 and Zolgensma for spinal muscular atrophy was the most expensive drug ever placed on the market with a price of $2.125 million [4]. This high cost is most likely due to the manufacturing workflow of these products, given that the entire manufacturing process is carried out outside Latin America. There is no doubt that in the long future this biotechnology tends to have a more accessible, as happened with BMT, but needs to be encouraged by the presence of qualified centers for manufacturing this biological products independently, outside from Europe or United States, as example as in Brazil.

In Brazil, the National Cellular Therapy Network (RNTC) has eight Cellular Technology Centers located in five Brazilian states and 52 laboratories selected by the CNPq (National Council for Scientific and Technological Development) and the Department of Science and Technology (Decit) of the Ministry of Health [5]. RNTC’s main objective is to increase integration between researchers from all over Brazil and facilitate the exchange of information about stem cell research carried out in the country, generating scientific knowledge and technological competence in regenerative medicine[5, 6]. RNTC is a partner of the International Society for Cellular and Gene Therapy (ISCGT) and the International Society for Stem Cell Research (ISSCR) to promote and disseminate cell and gene therapy (CGT) knowledge. Their aim is to support CGT in Brazil and allow the development of good manufacturing practices (GMP) locally as a source for cell therapies products [4]. To try and accelerate this type of initiatives, the Brazilian Ministry of Health created, in 2009, the Program to Support and Development Unified Health System (PROADI-SUS). Through this alliance between six reference hospitals in Brazil and the Ministry of Health, activities of human resources training, research, evaluation and incorporation of technologies, management and specialized assistance projects required by the Ministry of Health are supported. In this scenario, one of the topics covered is advancing cell therapy [7].

Brazilian researchers are working hard to develop national cell and gene platforms to produce these therapies products *in loco*. This will certainly contribute to minimize the costs and offer this promising therapy more quickly for Brazilian patients. In this context, our group proposed to construct a natural killer (NK) cell expansion platform.

NK cells are being increasingly explored in the field of cell therapy due to their innate antitumor characteristics. NK cells have a very singular mechanism for distinguishing between tumor and healthy cells, based on the missing-self theory – a balance with NK group-2 receptors (NKG2 type) and killer-cell immunoglobulin-like receptors (KIRs) [8, 9]. Because of these characteristics, this specific immune cell is considered a potential off-the-shelf cell therapy[10].

Hematopoietic stem cell transplantation (HSCT) combined with NK cell adoptive immunotherapy results in better engraftment and positively modulates the graft-versus-host response, reducing the risk of relapses and favoring remission in patients with leukemia [9, 11]. In addition, studies demonstrated that when associated with NK cells therapy, chemotherapy shows promising results for the treatment of solid tumors such as glioblastoma [12, 13].

Despite NK cells therapy being quite promising, the amount of NK cells necessary for adoptive immunotherapy is not available from natural resources as peripheral blood (5%-15%) [11] and umbilical cord blood (expression around 30%) [14]. The ideal number of cells for adoptive immunotherapy is 1×10^5^ to 1×10^9^ NK cells per kilogram of patient’s body weight. This is the amount that makes it possible to achieve a therapeutic response [15, 16]. In this context, the development of feeder cells to *ex vivo* NK cell expansion is crucial for the broad application of this therapy.

Feeder cells are genetically modified to express specific proteins capable to expand, activate, and increase the cytotoxicity of NK cells due to insertion exogenous proteins by lentivirus. The most common lineage used is the K562 cells line transduced to express 4-1BBL (CD137L, tumor necrosis factor receptor superfamily 9 ligand), with the functional portion of interleukin (IL)-21 in their membrane (mbIL-21, membrane bound interleukin 21). Together, they stimulate the proliferation and activation of NK cells without induction of early senescence [17–19].

The co-cultivation with feeder cell and NK cell is essential to developing this cell therapy. However, although this concept is well established, the centers that develop their feeder cells do not share their products due to intellectual property issues (patents). Inspired by Ojo et al. (2019), a group who produced their owner feeder cell, we aimed, in this work, to construct a national feeder cell from the K562 cell transduced with mbIL-21 and 4-1BBL that can promote NK cells proliferation with cytotoxicity preserving, and without inducing senescence or exhaustion.

## MATERIAL AND METHODS

### Isolation of human NK cells and ethical aspects

Peripheral blood was collected from five healthy donors to obtain NK cells, and from one patient with acute myeloid leukemia to acquire blasts for the cytotoxicity test. The collection occurred at the Hospital Israelita Albert Einstein from August 11, 2021 to March 9, 2022. All procedures performed in studies involving human participants were in accordance with the 1964 Helsinki Declaration and its later amendments. The Research Ethics Committee of the Hospital Israelita Albert Einstein approved this study under protocol number 06191819.9.0000.0071. All donors provided informed written consent to be part of the study, and their personal details were removed from this analysis.

### Production of lentiviral vectors containing mIL21 or 4-1BBL

The lentiviral vectors were produced using HEK293 T expression system. Briefly, HEK293T cells were transfected with packing plasmid psPAX2 (addgene, 12260), envelope plasmid pmD2.G (addgene,12259) and plasmids carrying on the sequence of mbIL-21 (Vector Builder, pLV and [Exp]-Puro-EF1A>Membrane Bound IL-21 [mbIL-21]) or expressing the sequence of 4-1BBL (Vector Builder, pLV[Exp]-Hygro-EF1A>4-1BBL) using X-tremeGENE™ HP DNA Transfection Reagent (Merck).

### K562.Clone1 feeder cell manufacture

K562 cells (CCL-243-ATCC) were obtained from Rio de Janeiro Cell Bank (BCRJ). The cell identity was confirmed through K562 STR DNA fingerprinting (https://www.atcc.org/products/ccl-243) by GenePrint 10 System Kit (Promega).

K562 feeder cells were transduced with lentiviral vectors containing genes of mbIL-21 or 4-1BBL and using 8μg/mL of Polybrene (Hexadimethrine bromide, Sigma-Aldrich), in accordance with Wald Lab protocol (Dr. David Wald - Case Western Reserve University, OHIO, USA) resulting in K562.Clone1, which expresses mbIL-21 and 4-1BBL on its surface.

All clones that expressed mbIL-21 and/or 4-1BBL had been previously selected with antibiotics; however, to improve and achieve high purity, K562.Clone1 was sorted at MoFlo Astrios EQ (Beckman Coulter). Feeder cells were expanded with RPMI1640 medium, 10% inactive bovine fetal serum (iBFS), 1% L-glutamine, 1% penicillin and streptomycin (Gibco, ThermoFischer). Before co-culture with peripheral blood mononuclear cells (PBMCs) or NK cells, K562.Clone1 cells were irradiated under 100 Gray (100cGy) in GamaCell3000 (Best Theratronics) [20, 21].

### NK cell expansion

Fresh blood cells were collected in ACD-A tubes (Sarsted) and mononuclear cells (MN) were isolated by Ficoll Paque Plus (GE Health). Next, PBMCs were co-cultured with irradiated K562.Clone1 in G-rex flasks (Wilson Wolf), using the ratio of 1:4 (PBMC: K562.Clone1 or PBMC: K562-WT), and 100 U/mL of IL-2 (Gibco) were added every two days.

On day 7, CD3 fraction (Miltenyi,130-050-101) were depleted by LD Collumn (Miltenyi Biotec), and the CD3-negative portions were co-cultured with irradiated K562.Clone1 or K562-WT (ratio 1:4) in G-rex flasks up to the 14th day. The cell proliferation was monitored by cell counting Neubauer chamber and immunophenotyping were performed on day 0, 7, and 14.

For control purposes, NK expansion was also performed with K562 without any modification (parental cell), here referred to as K562-WT.

### Characterization of *ex vivo* expanded NK cells

*Ex vivo* NK cells were characterized by different markers as shown in **Table 1**. For immunophenotyping, the cells were incubated with monoclonal antibodies for 30 minutes and then washed twice with FACS Buffer (PBS+0,1%BSFi). NK cells were characterized using CD45 (clone HI30, BD), CD56 (clone NCAM16.2, BD) and CD16 (clone 3G8, BD), Live/Dead Fixable Aqua Dead Cell Viability Reagent (405 nm excitation – Invitrogen - L34957), anti-TCR-Ɣδ (clone B1, BD), and anti-CD3 (clone SK7, BD) to exclude unwanted cells. NK activator and inhibitory receptors, such as NKG2C (clone 134591, BD), NKG2D (clone 1D11,BD), DNAM-1 (clone 11A8,BD), NKp30 (clone 210845, R&D System), NKp44 (clone P44-8, BD) NKG2A (clone 131411, BD), KIR3DL1 (clone DX9, Biolegend), KIR2DL2/3 (clone DX27, Biolegend), and KIR2DL1 (clone HP-MA4, Biolegend) were also evaluated.

**Table 1.**
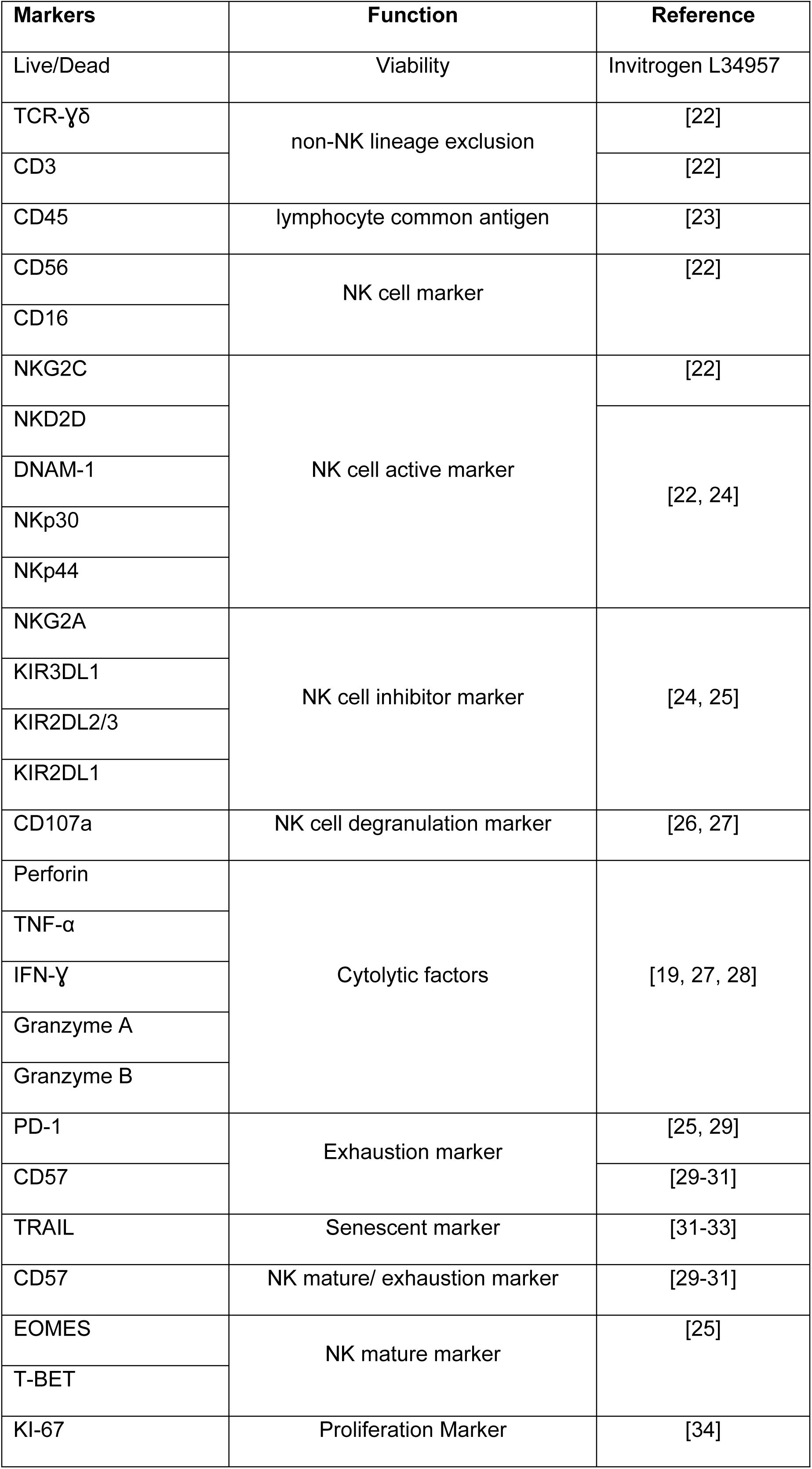
Characterization of natural killer cells.

Degranulation, senescence, and exhaustion of NK cells were assessed. Intracellular stained were required and NK cells were labeled and incubated with antibodies and FIX & PERM Cell Permeabilization reagent (Invitrogen), as described by manufacturer.

The cytotoxic markers included CD107a (clone H4A3, Biolegend), perforin (clone δG9,BD) TNF-α (clone MAB11, Biolegend), IFN-Ɣ (clone B27,BD), granzyme A (clone CB9,Biologend), and granzyme B (GB11, BD). For senescence and exhaustion NK cell evaluation the following antibodies were used: anti-PD-1 (clone EH12.2H7, Biolegend), - CD57 (clone NK-1, BD), -TRAIL (clone RIK-2, BD), -TIM-3 (Clone 7D3, BD), -KI-67 (clone B56, BD), EOMES (clone WD1928, Ebioscience/Thermo Fisher), and T-BET (clone 4B10, Biolegend).

Cells were acquired using BD LSRFortessa (BD Biosciences) or on Attune NxT (Thermo Fisher Scientific) flow cytometry. The analysis was performed using Flowjo 10 software (BD).

### Cytotoxicity assay

NK cell cytotoxic function was assessed by Calcein-AM (CAM) assay. The target tumor cells were K562 (human erythroleukemia lineage), OCI-AML2 (acute myeloid leukemia lineage), AML Blast (from a patient diagnosed with AML), and U-87 MG (glioblastoma lineage), the last kindly provided by Dr Keith Okamoto. Briefly, target cells were labeled with 25nM of Calcein-AM for 30 minutes, then washed twice with phosphate buffered saline (PBS) + 5% SFB (s protein flob-based assay). Next, cells were resuspended to 10^4^/ml in complete medium.

Labeled target cells were placed in 96-well Round Bottom Microwell Plates (Corning), with NK:T ratios ranging from 0.5:1 to 40:1, in triplicate. The controls were: non stained target cells, CAM-stained cells and labeled target cells in medium plus 2% Triton X-100. The prepared assay was incubated at 37°C in 5% CO_2_ for 4 hours. Samples were acquired in LSR Fortessa (BD) or Attune (Thermo Fisher).

### Statistical analysis

Statistical analysis was performed using Prism 8 software (Graphpad). The data were presented as means and standard deviations (range, minimum-maximum). The difference between groups was determined by 2-way analysis of variance (ANOVA), or 3-way ANOVA. Multiple t-test was used to evaluate ratios or markers. Wilcoxon signed-rank test was used to compare the median of samples, considering hypothetical value = 0 used in the calcein assay. We considered as statistically significant p values of 0.05, and we also give denotations for comparisons where p<0.01 and p<0.001.

## RESULTS

### NK cells fold expansion

The mean number of PBMCs seeded in the co-culture of the five donors blood was 1.80 ± 0.45 × 10^6^ cells, and the mean percentage of NK cells was 5.95% ± 3.92% (range, 2.1 - 12.1%) on day 0. From day 0 to day 7, an average of 3.23 ± 2.19 × 10^6^ mononuclear cells (MN) (range 1.30 - 6.26× 10^6^; p<0.0001) was found and 60.77% ± 9.76% CD3^−^CD56^+^ cells expanded with K562.Clone1. When MN cells were cultivated with K562-WT, there was a presence of 2.72 ± 1.45 × 10^6^ cells (range, 1.36 - 5.00× 10^6^) and of 64.90% ± 18.53% CD3^−^CD56^+^ cells (range 51.8 - 78%) (**Figure 1A**). The highest numbers were obtained at day 14, when NK cells expanded in co-culture, with K562.Clone1 achieving 179.22 ± 75.70 × 10^6^ cells (range, 131.4-308.25× 10^6^), with an average of 83.97% ± 8.19% CD3^−^CD56^+^ cells (range, 78% - 93.3%) — compared to the co-culture with K562-WT that achieved 17.58 ± 10.34 × 10^6^ cells (range, 9.60 - 34.05× 10^6^) with an average of 75.15% ± 7.57% CD3^−^CD56^+^ cells (range 69.8% - 80.5%; p<0.0001, 2-way ANOVA) (**Figure 1A**). Thus, NK cells co-cultured with K562.Clone1 expanded more than 100 folds compared to NK cells co-cultivated with K562-WT (less than 10 folds) (**Figure 1B**) (p<0.0001, 2-way ANOVA).

**Figure 1.**
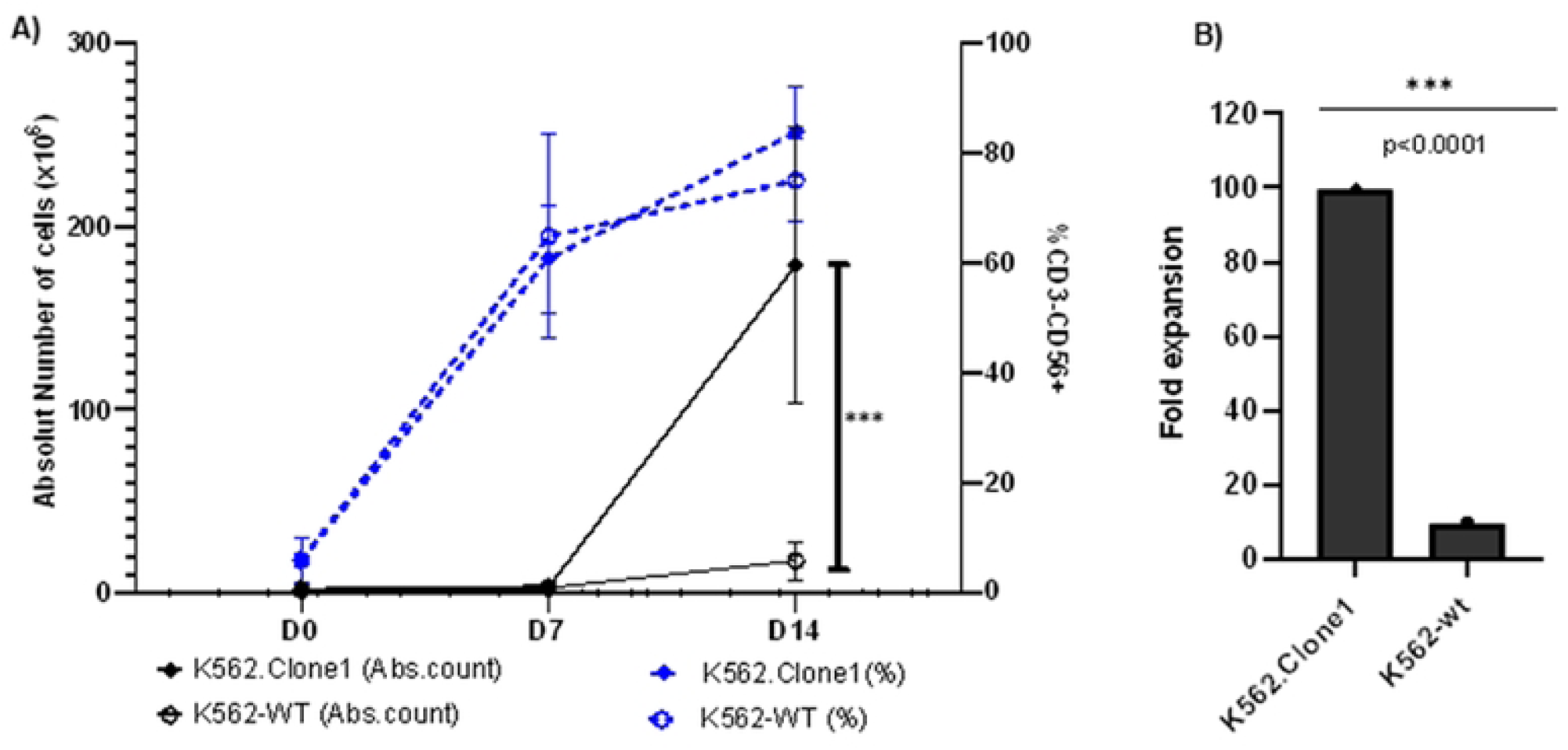
Natural killer (NK) cells derived from peripheral blood mononuclear cells (PBMC) from five blood donors, cultivated with K562.Clone1 or K562-WT. **A)** Black line: absolute number of cells on day O(DO), day 7 (D7) and day 14 (D14). Blue line: average of percentage NK cells (CD3^−^ CD56+) expression on day 0 (D0), day 7 (D7) and day 14 (D14) for both co-cultures. **B)** Fold expansion of NK cells after 14 days co-cultured with K562.Clone1 - expansion of ∼100 folds versus co-cultured with K562-WT - expansion of ∼10 folds (p<0.0001, 2-way ANOVA).

### NK cell cytotoxic evaluation

The NK cell cytotoxicity markers were measured on day 14. To perform the analyses, NK cells expanded with K562.Clone1 or K562-WT were subdivided into NK^dim^ (CD45+CD3-CD56+CD16+) and NK^bright^ (CD45+CD3-CD56+CD16-), the percentual cytotoxicity markers distribution was very similar in both co-cultures, no significant statistic was found. The median, minimum and maximum values were described in **Table 2** and represented in **Figure 2 A, B**.

**Figure 2.**
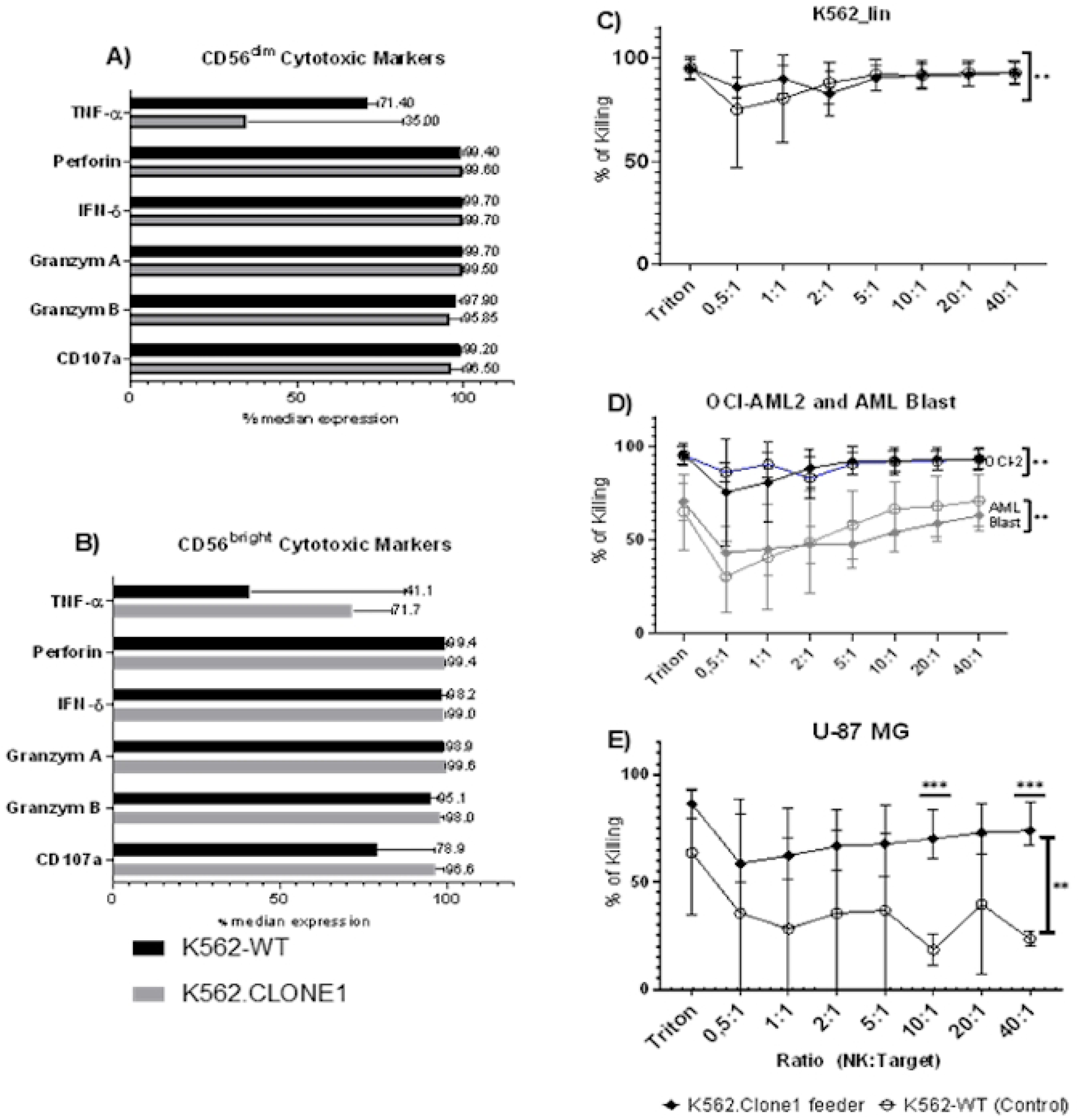
Evaluation of cytotoxic markers and functional assay in natural killer (NK) cells. Median of TNF-α, Perforin, IFN-Ɣ, Granzyme A, Granzyme B, and CD107a in NK^dim^**(A)** and NK^high^ **(B)** subpopulation, co-cultured with K562-WT (in black bar) and K562.Clone1 (in grey bar). No significant statistical was found between NK subtypes or co-cultures. **C - D)** Killing ratio for NK expanded with K562.Clone1 (♦) and K562-WT (◯) against leukemia cell lineages **E)** NK cells cytotoxic potency against glioblastoma lineage (U-86MG). The NK cells expanded with K562.Clone1 showed more cytotoxicity thanwhich expanded withK562-WT. The most important proportions were 10:1 and 40:1 (p<0.0001, multiple t-test)

**Table 2.** Cytotoxic markers distribution (medians and ranges) in natural killer cells co-cultured with K562.Clone1 and K562-WT.

|  | NK - CD56<sup>dim</sup> |  | NK - CD56<sup>bright</sup> |  |
| --- | --- | --- | --- | --- |
| Co-cultured with | K562.Clone1 | K562-WT | K562.Clone1 | K562-WT |
| **TNF-α** | 71.40% (range<br>54.30%-74.40%) | 35.00% (range<br>21.00%-82.10%) | 71.70% (range<br>53.00%-83.40%) | 41.10% (range<br>27.30%-87.10%) |
| **Perforin** | 99.40% (range<br>76.60%-100.00%) | 99.60 % (range<br>97.50%-100.00%) | 99.40% (range<br>77.10%-99.80%) | 99.40 % (range<br>97.30%-100.00%) |
| **INF-γ** | 99.70% (range<br>97.10%-99.90%) | 99.70 % (range<br>92.10%-100.00%) | 99.00% (range<br>93.10%-99.70%) | 98.20% (range<br>81.00%-100.00%) |
| **Granzym A** | 99.70% (range<br>98.80% -100.00%) | 99.50% (range<br>94.10%-99.80%) | 99.60% (range<br>98.30%-99.80%) | 98.90% (range<br>92.3%0-99.10%) |
| **Granzym B** | 97.90% (range<br>96.50%-99.60%) | 95.85% (range<br>92.50%-99.70%) | 98.00% (range<br>96.80%-99.20%) | 95.10% (range<br>91.40%-97.10%) |
| **CD107a** | 99.20% (range<br>95.40%-99.50%) | 96.50% (range<br>85.20%-100.00%) | 96.60% (range<br>90.90%-99.20%) | 78.90 % (range<br>63.10%-96.50%) |

The measurement of the cytotoxicity potency and efficiency against chronic myeloid leukemia lineage (K562), blasts from AML patient, AML lineage (OCI-AML2) and glioblastoma (U-86MG) were evaluated after the 14 days culture, using the calcein-AM assay. Samples with triton were considered positive controls.

The NK cells expanded with K562.Clone1 feeder were able to kill 91.09% (range, 82.97% - 92.99%) of K562, 91.96% (range, 75.32% - 93.00%) of OCI-AML2, and 50.76% (range, 43.03% - 62.84%) of the AML blast from a patient with AML (p>0.001, for all cell types). Similar ratios were seen in NK cells expanded with K562-WT, exhibiting similar median of cytotoxicity of 91.96% for K562 (range 75.32% - 92.95%), 91.09% for OCI-AML2 (range, 82.97% - 95.00%), and 61.42% for AML blast (range, 30.28%-70.73%) (**Figure 2 C, D**) (p>0.001, Wilcoxon test for all cell types). For U-86MG, NK cells expanded with K562.Clone1 cells presented a cytotoxicity median of 68,90% (range, 58.60% - 73.80%), while NK cells expanded with K562-WT showed a cytotoxicity median of 35.40% (range, 18.49% - 39.53%) (p<0.005, Wilcoxon test). The most significant killing ratio was 10:1 (effector cells:target cells) and 40:1 (p<0.001, multiple t-test) (**Figure 2E**).

The T-sNE graph in **Figure 3 A-H** shows cytotoxicity data in more detail, analyzing the mean fluorescence intensity (MIF) distribution of CD56, CD16, TNF-α, perforin, IFN-Ɣ, granzyme A, granzyme B and CD107a. By comparing each graph marker, it is possible to observe more hotspots in NK cells expanded with K562.Clone1 than with the controls (with a statistically significant difference compared with the MIF markers and co-culture; p<0.0001, 2-way ANOVA). There was a homogeneous dimensional distribution for NK cells co-culture with K562.Clone1, as a main cluster expresses high density of CD56 and CD16 expression, differently from NK cells expanded with K562-WT, which showed three others clusters around the main one (**Figure 3 A,B**).

**Figure 3.**
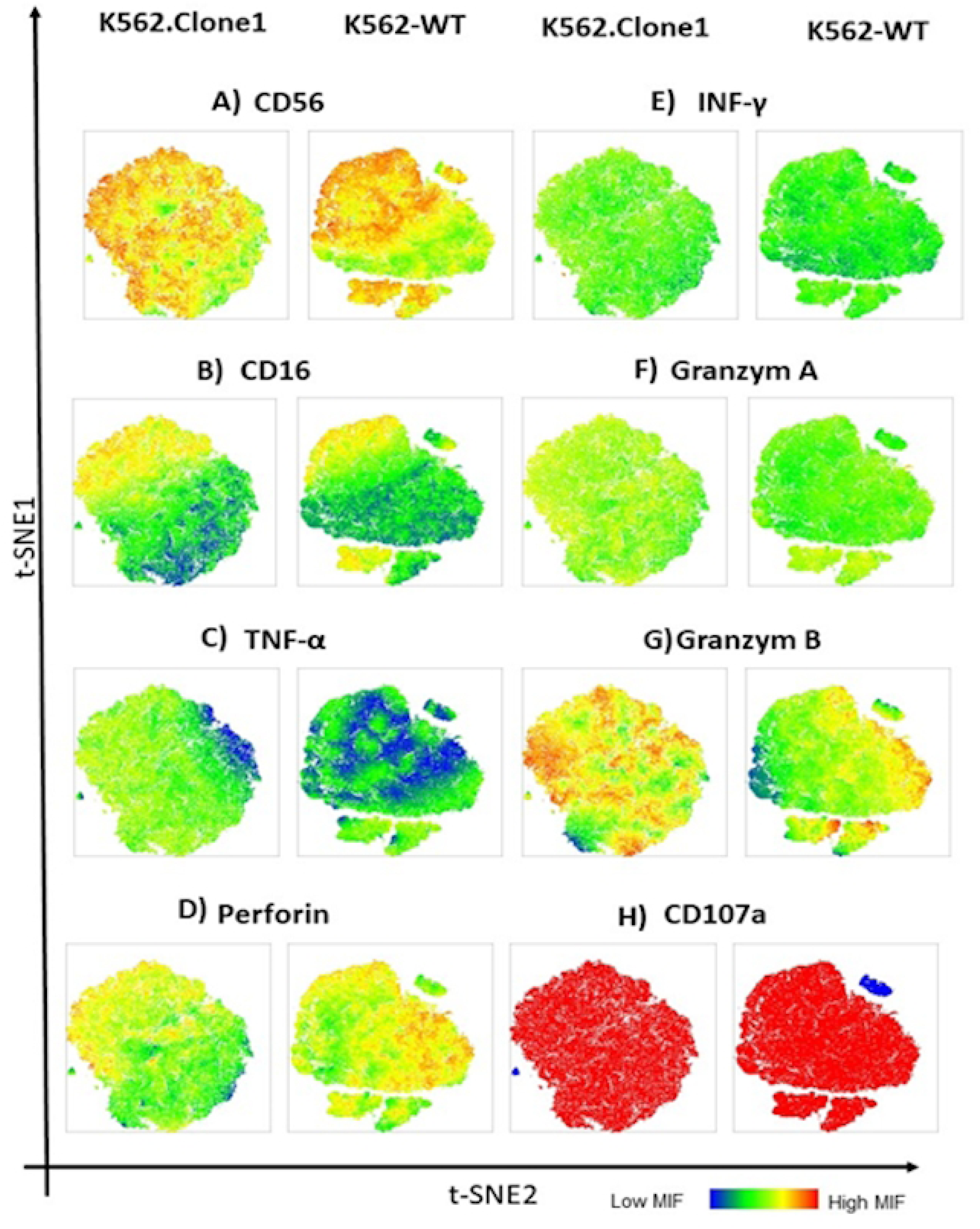
T-sNE graph for CD56, CD16, TNF-α, perforin, IFN-Ɣ, granzyme A, granzyme B and CD107a markers from natural killer (NK) cells expanded with K562.Clone1 (left) and K562-WT (right). The majority high MIF are present in the NK cell expanded with K562.Clone 1, except in perforin (D). Statistical analyses showed significant differences toward the markers and type of co­culture performed (K562-Clone1 or K562-WT). NK cells expanded with K562-WT showed central cluster with additional 3 clusters, which could be related with absence interaction with feeder cell ligands (mbll-21 and 4-1BBL). MIF for CD16 (B), perforin (D), INF-g(E) granzyme A (F) and CD107a (H), were very similar. MIF for CD56 (A), TNF-a (C) and granzyme 8 (G) were high in NK co-culture with K562.Clone1. Heat map scale: low MIF ≤300 to high MIF ≥11,764.2.

### NK cell functionality

The mean percentage and SD of activator markers (NKG2C, NK2G2D, NKp44, NKp30, DNAM-1) and inhibitory markers (NKG2A, KIR3DL1, KIR2DL2/3, and KIR2DL1 of NK^dim^ and NK^bright^ cultivated with K562.Clone1 or K562-WT (**Table 3**). The table shows that there is a higher expression of activation than of inhibition markers, except for KIR3DL1 and NKG2A, which were highly expressed in all subtypes of NK, independently of the type of co-culture used.

**Table 3.** Expression of activation and inhibition markers in natural killer CD56^dim^ and CD56^bright^ cells co-cultured with K562.Clone1 and K562-WT.

|  |  | NK - CD56<sup>dim</sup> |  | NK - CD56<sup>bright</sup> |  |
| --- | --- | --- | --- | --- | --- |
| Co-cultured with |  | K562.Clone1 | K562-WT | K562.Clone1 | K562-WT |
| NK activation markers | NKG2C | 85.00% ± 11.96% | 97.85% ± 1.77% | 82.08% ± 12.56% | 96.35% ± 1.77% |
|  | NKG2D | 99.68% ± 0.46% | 100%±0 | 99.94% ± 0.05% | 100%±0 |
|  | NKp44 | 99.84% ± 0.25% | 100%±0 | 99.92% ± 0.08% | 100%±0 |
|  | NKp30 | 99.68% ± 0.04% | 100%±0 | 99.98% ± 0.04% | 100%±0 |
|  | DNAM-1 | 0.41% ± 0.79% | 100%±0 | 0.10% ± 0.20% | 100%±0 |
| NK inhibition markers | KIR2DL1 | 32.10% ± 28.65 | 22.10%± 9.76% | 57.74% ± 31.44% | 45.05%± 1.89% |
|  | KIR2DL2/3 | 32.98% ± 13.07% | 28.00%± 3.39% | 63.04% ± 11.15% | 57.55% ± 5.30% |
|  | KIR3DL1 | 90.68% ± 19.72% | 98.85%± 0.07% | 100%±0 | 100%±0 |
|  | NKG2A | 78.98% ± 17.22% | 85.10% | 86.14 ± 12.27% | 76.37 ± 23.84% |

**Figure 4A** presents the distribution of these values graphically, confirming the activation signals as predominant. For example, statistical analyses comparing NK^dim^ or NK^bright^ with their respective controls cultivated with K562.Clone1 or K562-WT have shown significant differences for activation (p < 0.0001, 2-way ANOVA) and inhibition (p < 0.01).

**Figure 4.**
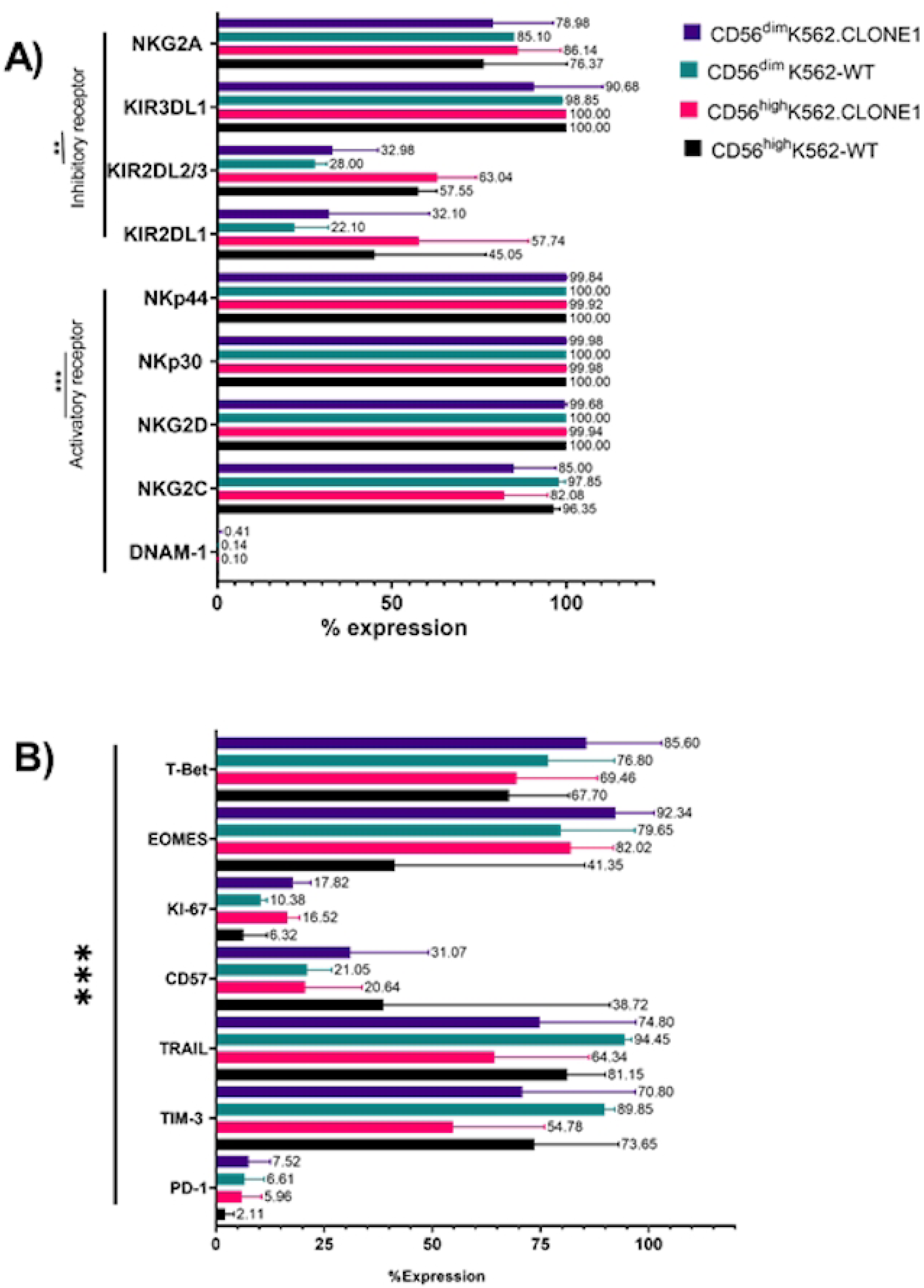
Activator and Inhibitor markers and NK cell balance. **A)** Percentual expression of activator NK cell markers (DNAM-1, NKG2C, NKG2D, NKp30, NKp44) and inhibitor markers (KIR2DL1, KIR2DL2/3, KIR2DL1, NKG2A) distributed according to NK cell subpopulation (dim and bright) and co-culture with K562.Clone1 or K562-WT. **B)** Major exhausted and senescent marker (PD-1) and associated markers (TIM-3, TRAIL, CD57, Kl-67, EOMES, T-BET) related with cytotoxic function, maturation, proliferation, and maturation to investigate the dysfunctional state.

We evaluated the state of anergy, senescence and cellular exhaustion carefully, considering the expression of cytotoxicity, activation, and inhibition markers. These dysfunctional conditions were defined by a set of markers such as: KI-67, involved in cell proliferation, CD57 and TIM-3, indicators of maturation stage, TRAIL (CD253), a ligand related to apoptosis induction, PD-1, a strong exhaustion marker, and transcription factors such as T-box transcription factor (T-bet), and eomesodermin (EOMES), associated with activation and inhibition markers. **Figure 5** shows the markers most related with functional, active and effector function and dysfunctional state.

**Figure 5.**
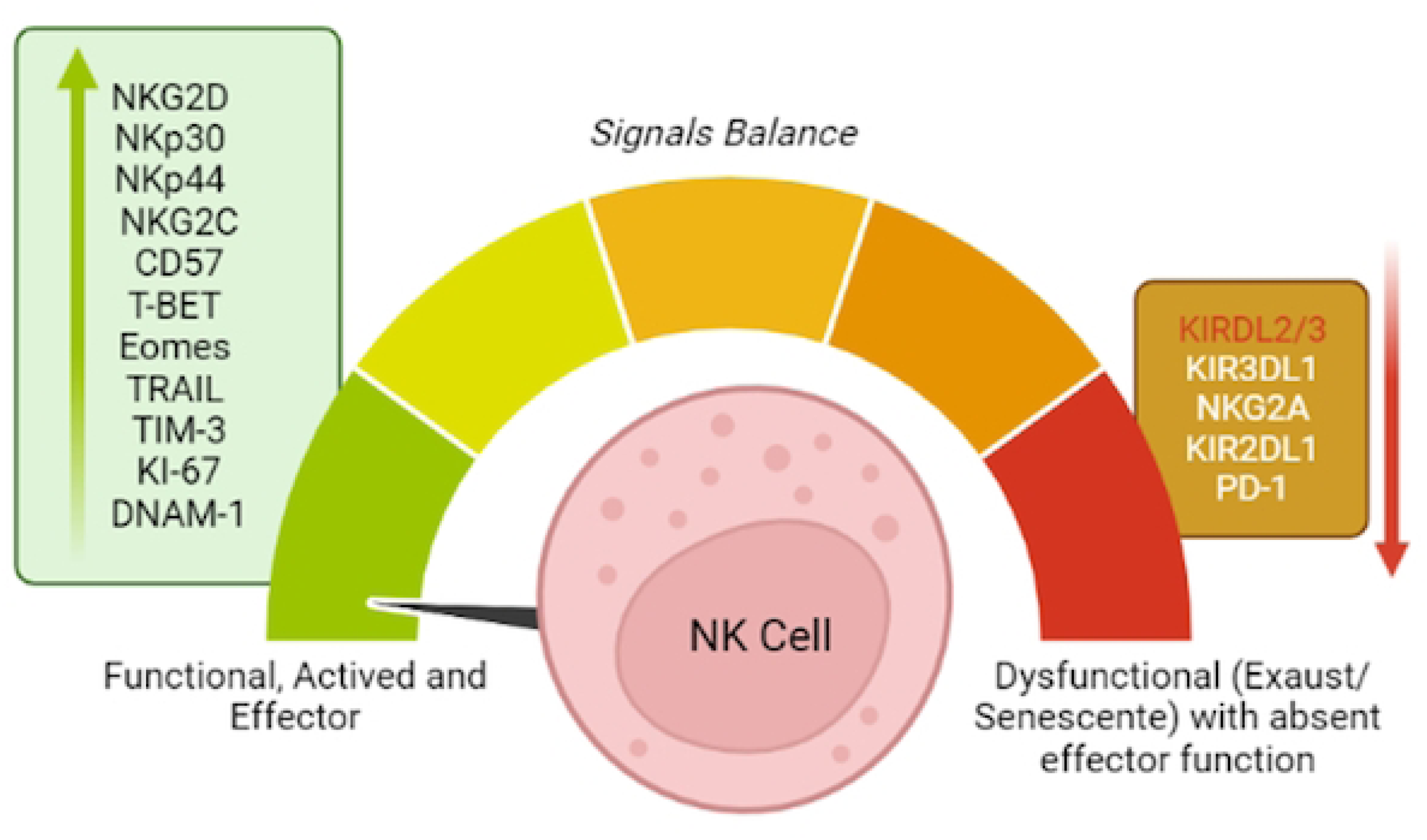
Key indicators of the cellular status of natural killer (NK) cells. On the left, the main markers of activation NK cell markers (DNAM-1, NKG2C, NKG2D, NKp30, NKp44), inhibition markers (KIR2DL1, KIR2DL2/3, KIR2DL1, NKG2A), exhausted and senescent marker (PD-1) and associated markers (TIM-3, TRAIL, CD57, Kl-67, EOMES, T-BET) related with thecytotoxic function, maturation, proliferation, and maturation, used to investigate the NK cells’ dysfunctional state.

In this way, PD-1, T-BET, EOMES, TRAIL, KI-67, CD57 and TIM-3 were evaluated in NK cells CD56^bright^ and CD56^dim^ expanded with K562.Clone 1 or with K562-WT. The means and SDs are described in **Table 4**, and the results are represented in **Figure 4B**.

**Table 4.** Evaluation of the senescent and exhaustion indicators expression in natural killer (NK) CD56^dim^ and CD56^bright^ cells co-cultured with K562.Clone1 and K562-WT.

|  | NK - CD56<sup>dim</sup> |  | NK - CD56<sup>bright</sup> |  |
| --- | --- | --- | --- | --- |
| Co-cultured with | K562.Clone1 | K562-WT | K562.Clone1 | K562-WT |
| PD-1 | 7.51% ± 5.03% | 6.60% ±4.58% | 5.96% ± 4.59% | 2.10% ± 2.08% |
| T-BET | 85.60% ± 17.37% | 76.80% ± 15.27% | 69.46% ± 18.70% | 67.70% ± 13.85% |
| EOMES | 92.34% ± 8.90% | 79.65% ± 17.18% | 82.02% ± 9.83% | 41.35% ± 43.91% |
| TRAIL | 74.80% ± 22.25% | 94.45% ± 1.62% | 64.34% ± 21.90% | 81.15% ± 8.83% |
| KI-67 | 17.82% ± 4.10% | 10.38% ± 1.44% | 16.52% ± 2.80% | 6.31% ± 5.49% |
| CD57 | 31.06% ± 18.07% | 21.05% ± 5.72% | 20.63% ± 13.11% | 38.72% ± 52.29% |
| TIM-3 | 70.80% ± 26.10% | 89.85% ± 2.33% | 54.78% ± 21.13% | 73.65% ± 19.44% |

Based on these data we verified that the low PD-1 expression associated with high expression of T-BET, EOMES and TRAIL showed the cytotoxic ability was conserved (**Table 4**). The expression of KI-67 means that NK cells co-cultivated with K562.Clone1 proliferates more quickly than those co-cultivated with K562-WT (**Table 4**). Moreover, NK cells seemed to be in mature stages, which could be inferred due to the presence of CD57 and TIM-3 markers. Statistical analyses were significant between CD56^bright^ and CD56^dim^, independently of the co-culture used (p<0.0001, 3-way ANOVA).

## DISCUSSION

NK cells have a natural cytotoxicity against tumor cells. They represent about 10% to 15% of the total peripheral blood lymphocytes in healthy individuals and about 25% to 30% of the lymphocytes present in cord blood (29, 31). Moreover, NK cells improve the clinical outcome in hematopoietic stem cell transplantation since they are associated to lower relapse rates and reduced incidence of graft versus host disease (GVHD) [2, 35, 36].

A phase I/II clinical trial conducted by Ciurea and colleagues (2022) demonstrated a positive association between AI of *ex vivo* expanded NK cells with feeder cells and clinical outcomes of AML patients undergoing HSCT. Patients were followed up for 24 months, and the relapse rate was 4% *versus* 38%, an index related to conventional treatment, not using AI. Furthermore, disease-free survival was 66% *vs*. 44% in cases and controls, respectively. All patients achieved first complete remission. Moreover, approximately 50% of AML patients included in the trial had grade 1 to 2 graft-versus-host disease (GVHD), with rare cases of grade 3 to 4, but no case of chronic GVHD was observed during the follow-up in patients treated with NK AI [15, 37]. The study demonstrated that infusion of higher doses of NK cells, ranging from 1×10^5^ cells/kg to 1 ×10^8^ cells/kg of body weight was associated with progressively higher numbers of NK cells at the beginning of post-transplantation. This finding suggests a dose-dependent effect, leading to an early immune reconstitution, avoiding relapse and longer disease-free time and very low death rate [37].

Another clinical study conducted by Otegbeye et al. (2022) evaluated the efficacy and safety of *ex vivo* expanded NK cells in patients with myelodysplastic syndrome (MDS), AML or colorectal cancer. The patients, previously submitted to chemotherapy with cyclophosphamide and fludarabine, received 2 doses of NK cells ranging from 1×10^7^ cells/Kg to 5 ×10^7^ cells/Kg. They were monitored for 100 days after AI. There was no occurrence of GVHD. The toxicity related to the doses of NK cells administered was limited, demonstrating the safety of using these cells in cancer patients [38].

Several clinical studies using NK cells are in progress, not only in their genuine forms, but also with chimeric modifications capable of promoting a targeted response. One example is CAR-NK [39], using Bi or Tri specific antibodies, which induces the death of tumor cells via CD16, through the process of cytotoxicity mediated by antibodies [40]. Another is using interleukin (IL) cocktails capable of inducing a memory in these cells, originating NK-memory-like cells, potent cells that have the ability to respond against lymphomas.[41]

In all mentioned studies the use of feeder cells was necessary to properly expand NK cells *ex vivo*. Vidard et al. (2019) observed that non-transduced K562 cells promote greater expansion in the first week, but due to the absence of 4-1BBL or mbIL-21, these cells do not have the ability to sustain a prolonged proliferative response [42]. Denman et al. (2012) showed that the use of this *ex vivo* expansion platform promotes the clonal expansion of NK cells, maintaining their viability, conserving and increasing their antitumor cytotoxicity [20, 43]. These feeder cells are co-cultured with NK cells, mainly from peripheral blood and umbilical cord blood, and they have the ability to expand these immune cells logarithmically, reaching the number of cells needed for therapeutic use.

In this context, we manufactured *in loco* feeder cells with successful transduction of mbIL-21 and 4-1BBL proteins, originating the K562.Clone1 cell. This feeder cell was capable to expand NK cells more than 70 times, conserving their phenotypical and cytotoxic features. Other associated molecules can be introduced in K562 to constitute a feeder cell. Fujisaki et al. (2009) produced a K562-mb15-41BBL feeder cell, which had the capacity to expand an average of 21.6 times the NK cells in the first 7 days.[44] Jiang et al. (2014), produced a K562.4-1BBL feeder cell capable to expand up to 1000 folds of NK cells after 21 days.[45] On the other hand, Denman et al. (2012) using feeder cells based on mbIL21, 4-1BB and CD86 achieved 31.74 to 47.96 folds of NK expansion after 21 days of co-culture[20]. In our work, the K562.Clone1 was able to expand up to 100 folds of NK cell, conserving their phenotypical features, cytotoxic function with no exhausting or senescent signals.

NK cells expanded *ex vivo* with the K562.Clone1 feeder cell expressed important markers related with their cytotoxic potentiality, such as CD107a, related with the degranulation ability of the NK cell. This protein is a glycosylated molecule that preserves the integrity of the lysosomal membrane and only degranulates when encounters a tumor cell [46]. Inside of these granules, cytolytic factors could be found, such as perforin and granzyme, responsible for direct lysing the tumor cell with induction of IFN-Ɣ and TNF-α, which induces cell death in tumor cells and switch on the modulation of the adaptive immune response [29, 47]. Additionally, the balance between activators and inhibitors signals are essential for the recognition and distinction of healthy or cancerous cells, [13, 48, 49].

Furthermore, NK cells expanded with K562.Clone1 show a higher expression of activator receptors as NKG2D, NKp30 and NKp44. NKG2D, a type II lectin family receptor, is a key marker for activator response in NK cells [46]. Usually, NKG2D ligands such as MIC-A, MIC-B, and ULBP-1, -2 and -3 are increased in carcinomas including melanoma, leukemia, myeloma, glioma, prostate, breast, lung, and colon, leading to the predominance of an activator response by NK cells [3, 50–52]. The interaction of NKG2D and its ligands triggers an activating response culminating in cytokine production, activation, and proliferation of NK cells. Other activating receptors, such as NKp30 and NKp44, also used this same signaling pathway, resulting in the movement of lysosomes containing factors such as perforin, INF-γ, and TNF-α. All these factors promote the elimination of tumor cells [26, 50, 53]. Fujisaki et al. (2009) observed a high expression of NKp30, NKp44, and intracellular factors granzyme B and perforin after treatment with feeder cells that carried the mbIL-21 protein [44].

On the other hand, we observed the expression of the inhibitory receptor KIR2DL2_3, which consist into two proteins that have approximately 94% homology: KIR2DL2 and KIR2DL3, highly polymorphic as well as their respective HLA-C ligand, leading to a polymorphism in the KIR receptor and HLA-C ligand which alters the avidity of binding, resulting in strong or tenuous inhibitory response [54]. The expression of inhibitory receptor and the interaction between KIR2DL1, KIR2DL3, KIR3DL1, and NKG2A with their respective MHC-class I ligands are very important since they are involved in the training of NK cells for the identification of self and non-self MHC. The complete absence interaction between inhibitory KIR and their ligands causes a hyperresponsiveness state of the NK cells similar state when the cells are senescent or exhausted [54–56].

In addition, cytotoxic data performed by the calcein AM assay supports the preservation of cytotoxic potency and function in NK cells expanded *ex vivo* with K562.Clone1. Our data show the high capacity of NK cells of killing leukemia tumoral lineage, blasts from AML patients, and glioblastoma. As shown by Kmiecik et al. (2013) and Golán et al. (2018), our cytotoxic assay exhibits the strong potential for NK cells of killing U86-MG, the glioblastoma lineage. Both studies indicated that NK cell therapy is promising in the treatment of glioblastoma or brain cancer, as the NK cells present in the meningeal region were associated with a better prognosis, due to CD16 activity via ADCC and TRAIL, via ligand and receptor interaction, leading to tumoral cell apoptosis [57, 58].

The interaction of NK cells and K562.Clone1 is capable of expanding NK cells in a robust way, without changes in their phenotype, without signs of senescence or exhaustion. Furthermore, NK cells presented high cytotoxicity in response to the positive expression of granzymes, perforin, INF-γ, TNF-α, and receptors such as CD16 and TRAIL (**Figure 6**).

**Figure 6.** Characteristics of natural killer (NK) cells expanded *ex vivo* with K562.Clone1 feeder cells. Multiparametric cytometry of *ex vivo* NK cells expanded with K562.Clone1 feeder cells was performed. NK cells present the classical markers, such as CD56, and also CD16, which is indispensable for antibody-dependent cellular cytotoxicity. In addition, the expression of TRAIL, activation KIRs (NKG2D and NKps), CD107a (a degranulation marker), associated with the expression of cytolytic factors such as granzyme A, granzyme B, INF, TNF and perforin, strongly indicates that the NK cells were activated and cytotoxic, and NK cells were able to kill tumor cells through these pathways. Moreover, the presence of inhibitory KIRs, as well the activation of KIRs suggest that these cells are capable to recognize and distinguish abnormal cells from healthy cells, a role played by the interaction between KlRs and their ligands by missing-self theory. K562.clone1 feeder cell, does not cause exhaustion conditions in NK cell after co-culture, this could be confirmed once PD-1 expression was very low. Moreover, NK cells expanded with K562.Clone1 expressed Kl-67, a major proliferation marker. In addition, we considered CD57 and TIM-3 proteins more related with cell maturation than with senescence or exhaustion in NK cells, given the high expression of T­ BET, EOMES, TRAIL and the other cytotoxic factors.

## CONCLUSION

Our group successfully built a national feeder cell, named K562.Clone1. The co-culture with K562.Clone1 feeder cell and peripheral NK cell sustains the activation and functionality of these NK cells, preserving their primordial functions such as missing-self, important to distinguish health and defective cell, and natural cytotoxicity against tumoral cells. This work offers Brazilian researchers a clinical platform for ex vivo expansion of NK cells in the future.

## ACKNOWLEDGMENTS

The authors are thankful to Lilian Tiemi Inoue (Centro de Ensino e Pesquisa Hospital Sírio Libanês) for supporting the cell authenticity profiling. Figure 5 and Figure 4C were created with BioRender.com. The authors also thank Patricia Logullo, PhD, CMPP (Palavra Impressa) for manuscript editing services.

